# Classification of gene signatures for their information value and functional redundancy

**DOI:** 10.1101/136499

**Authors:** Laura Cantini, Laurence Calzone, Loredana Martignetti, Mattias Rydenfelt, Nils Blüthgen, Emmanuel Barillot, Andrei Zinovyev

## Abstract

Large collections of gene signatures play a pivotal role in interpreting results of omics data analysis but suffer from compositional (large overlap) and functional (redundant read-outs) redundancy, and many gene signatures rarely pop-up in statistical tests. Based on pan-cancer data analysis, here we define a restricted set of 962 so called informative signatures and demonstrate that they have more chances to appear highly enriched in cancer biology studies. We show that the majority of informative signatures conserve their weights for the composing genes (eigengenes) from one cancer type to another. We construct InfoSigMap, an interactive online map showing the structure of compositional and functional redundancies between informative signatures and charting the territories of biological functions accessible through transcriptomic studies. InfoSigMap can be used to visualize in one insightful picture the results of comparative omics data analyses and suggests reconsidering existing annotations of certain reference gene set groups.

## INTRODUCTION

The majority of the studies exploring gene expression data result in one or more gene signatures, i.e. list of genes sharing a common pattern of expression that can be employed to classify an independent dataset. Together with such “data-derived” signatures, “*a priori* knowledge-based” gene signatures are produced from the available gene ontologies or pathway databases. In recent years, data-derived and *a priori*-knowledge-based reference gene signatures have been widely employed to interpret the results of gene expression data analyses (e.g. differential expression, clustering). The number of available signatures is getting larger allowing users to benefit for a more exhaustive coverage of the existing biological processes. However, not all the signatures contained in these compendia are equally informative and the number of gene sets representing the same biological process is not equilibrated. Intrinsic redundancy and the presence of numerous signatures which have small chances to be selected in any analysis affects the results by heavy p-value corrections producing a higher number of false negative results. Conceptually, the aforementioned gene set redundancy can be of two types: compositional or functional (see Figure 1A). Compositionally redundant signatures are characterized by a large intersection in terms of the composing genes. On the opposite, two signatures will be here called functionally redundant when they represent two different (sometimes, with zero overlap) transcriptional read-outs of the same biological process. The presence of multiple functionally redundant signatures affects the enrichment analysis by highly scoring multiple gene sets belonging to analogous or related biological processes hiding other potentially relevant hits. Of note, any estimation of the functional redundancy is conditioned on the context (e.g., certain cancer type) and, therefore, depends on the data corpus which defines functional (e.g., correlation-based) relations between the individual genes.

**Figure 1.**
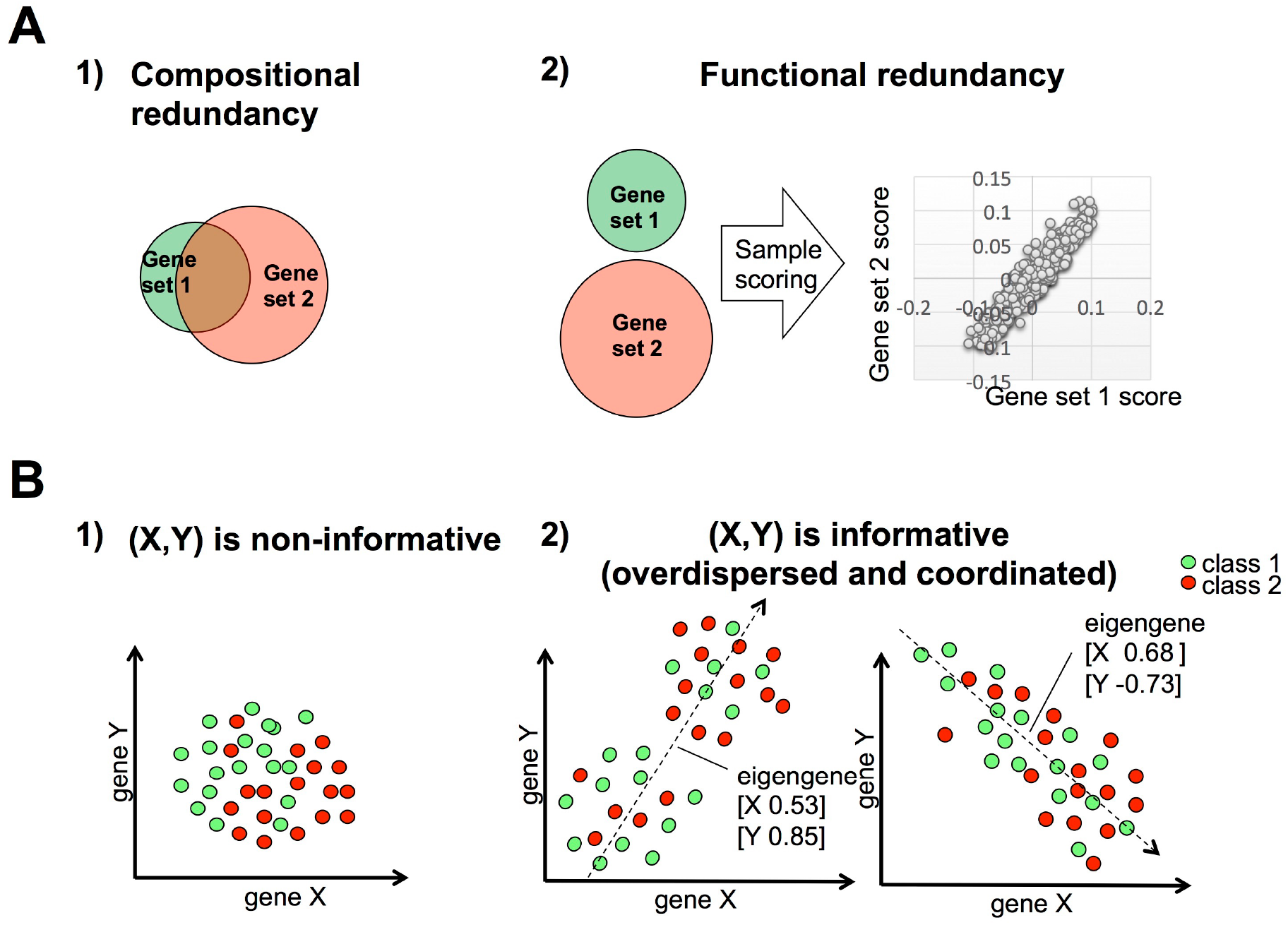
Schematic explanation of the basic notions used in this study. Panel A) schematically summarizes the two possible forms of redundancy between two gene sets. 1) Compositional redundancy corresponds to gene set overlap. 2) Functional redundancy represents instead different transcriptional read-outs of the same biological process and it is possible even for the gene sets with no overlap. Measuring functional redundancy depends on the way a sample is scored based on the expression of its genes and the chosen corpus of data. Panel B) explains the difference between non-informative (1) and informative (2) gene sets. A signature composed of two genes X and Y is here considered. The circles denote biological samples and the two colors correspond to two different labels : class 1 and class 2 (e.g., metastatic vs. primary tumors). Scatter plots are used to represent the expression values of gene X (X-axes) and gene Y (Y-axes) in each sample. Three types of samples distributions are shown. In 1) (isotropic case) no naturally distinguished axis in the points distribution, labeling of samples is needed to define their ranking. In 2) instead, it exists a distinguishable axis in the data distribution that allows a robust ranking of the samples independently on their labeling. Both second and third scenario leads to overdispersion and coordination of the corresponding gene set and are selected in the analysis.

To our knowledge, few methods have been proposed to address the problem of gene signature redundancy ^1–6^. Currently, the best attempt to define a robust and not-redundant collection of signatures is represented by the MSigDB Hallmarks (H) collection ^7^. H was obtained by merging and re-organizing compositionally redundant signatures and then refining the genes of the resulting signatures based on their ability to discriminate the associated phenotype. The Hallmarks collection methodology thus involves a manual curation and redefinition of signatures steps, which might create certain bias *vis a vis* an expert’s opinion leading to the loss of certain signature properties. More importantly, H, as all the other currently proposed procedures, takes into account only compositional redundancy without exploiting the problem of the functional one.

In this paper, a new approach to prioritize and classify gene signatures is proposed. Our method is based on the concept of “an informative signature”, which is capable of defining a ranking of samples independently on their labeling. Considering the simplest case of a gene signature composed of only two genes X and Y, their co-variance can define three possible scenarios of samples distribution (ranking), as reported in Figure 1B. Whereas the ranking defined by informative signatures presents a distinguishable axis (anisotropy), labels are required to define a samples ranking in the non-informative ones. As a consequence, when the dysregulation of an informative signature is tested on a two-conditions transcriptomics dataset, a significant enrichment score will be observed whenever the samples membership matches the direction of the largest variance and this is significantly greater than randomly expected (the gene set is “overdispersed”). In all other cases, a very specific distribution of sample labels is needed to obtain a significant enrichment score. Therefore, informative signatures, systematically defining robust and “objective” sample ranking in many datasets, are more valuable for data analysis.

Starting from a vast collection of signature compendia, composed of 12096 *a priori*-knowledge and data-derived signatures, we defined a restricted collection of 962 informative signatures, which is made available to the users for further applications. The collection was defined by exploiting a large pan-cancer TCGA collection of transcriptomic profiles (32 cancer types with totally 8991 transcriptomic profiles) and it is thus cancer biology-oriented. Of note is the fact that among the databases under investigation, a relatively small SPEED collection ^8^ proved to be the most informative with 55% of its signatures being highly informative. The reliability of our signature collection was then validated by comparing their performances with those of the starting complete collection in some typical data analysis scenarios. In all the considered examples, the informative gene sets were found much more frequently significant than the others, confirming the rationale behind the selection procedure here proposed.

Each signature defines a set of weights for the composing genes (which we term eigengene ^9^) in each analyzed dataset. We introduce the notion of “conserved signatures”, i.e. those signatures whose eigengenes are highly correlated across different cancer types. 73% of our informative signatures resulted to be also highly conserved, pointing out that they map well some universal functional blocks of the cellular machinery. The collection of informative gene signatures was classified by computing the average correlation between the sample activity profiles of all signatures across 32 cancers types. This metrics was used as a measure of functional redundancy in our analysis. We found that functional redundancy is an abundant phenomenon that does not result from a significant intersection size of two gene sets. Therefore the Hallmarks and the other previous works, employing only the overlap as a measure of redundancy, are capturing only a small portion of this phenomenon. In order to visually and interactively represent the structure of functional redundancies between informative gene signatures, we developed InfoSigMap (http://navicell.curie.fr/pages/maps_avcorrmodulenet.html), a user-friendly interactive Google Maps-based tool whose nodes are constituted by our set of informative signatures and whose links represent the two types of redundancies (compositional and functional). InfoSigMap can be used as a data visualization tool to provide a quick navigation into any set of scores associated to the informative signatures (e.g., enrichment scores). The use of InfoSigMap is demonstrated in some typical data analysis scenarios showing that it is able to provide a concise and biologically meaningful holistic view on the pattern of differential regulation of various cellular functions.

## RESULTS

### Informative signatures represent a small fraction of the widely employed gene sets

Defined the concept of “an informative signature” (see Methods), a large pan-cancer TCGA compendium of gene expression data derived from 32 solid cancer types was employed to restrict the input collection of gene signatures (12096) to 962 (see Supplementary Table S2) informative for cancer data analysis and corresponding to our compendium (for the selection procedure and inputs see Methods). Of the 962 identified informative signatures the majority were data-derived (231 knowledge-based, 706 data-derived, 15 MSigDB Hallmark and 10 MSigDB C1), showing that for cancer-oriented applications data-derived signatures tend to be more informative than knowledge-based ones. In order to assess which of the input compendia was more informative, the ratio between the number of informative gene sets and the total number of contained signatures was evaluated (Table 1). As shown in Table 1, the most informative compendium resulted to be SPEED (55% of informative signatures). The reliability of this database is thus corroborated by our results, suggesting that when dealing with cancer transcriptomics data analysis it should be probably preferred to alternative ones. Overall, good performances were also obtained by CIT (28%), the MSigDB C4 (31%) and the Hallmarks (H) (30%). While the best performing knowledge-based database resulted to be ACSN, with 13% of informative signatures. The distribution of the number of tumors in which the informative signatures were found to have the ROMA L1 and L1/L2 p-values significant were then investigated. As shown in Figure 2A and Supplementary Table S2, the majority of the informative signatures is cancer-specific (significant in only 2 cancer types), another peak is present around 15-20 tumor types and only 8 signatures are pan-cancer significant (significant in more than 25 cancer types). Furthermore, data-driven signatures tend to be more frequently pan-cancer-wise informative than the knowledge-based ones.

**Figure 2.**
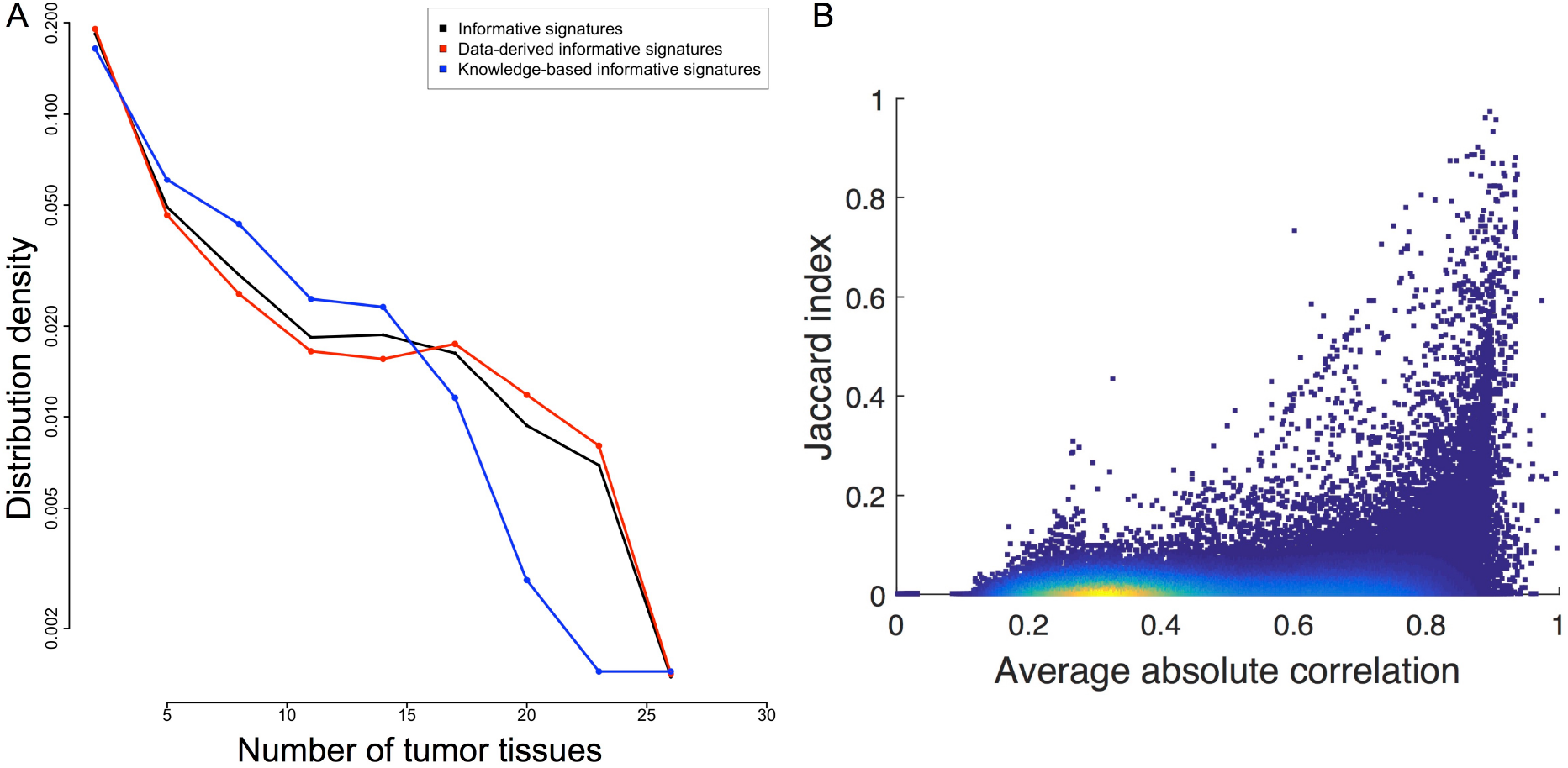
Behavior of the informative signatures across the different cancer transcriptomes. The main results regarding the pan-cancer behavior of the informative signatures are here summarized. In panel A) the distribution of the numbers of cancer types in which the informative signatures have ROMA L1 and L1/L2 p-values significant is reported. The behavior of all informative signatures is represented in black, that of data-driven and knowledge-based informative signatures is denoted in red and blue, respectively. In B) the dependence between informative gene sets overlap (Jaccard-index) and average correlation between the meta-samples defined by the informative gene sets is reported. Each point corresponds to an informative signature and their color is proportional to the points density : from red (high density) to blue (low density).

**Table 1.** Contribution of each signature collection to the informative set. Number of informative signature, total dimension and the fraction of the previous two fields are reported for each signature collection.

| Signatures collections | Data-Derived |  |  |  |  |  | Knowledge-based |  |  |  | H | C1 |
| --- | --- | --- | --- | --- | --- | --- | --- | --- | --- | --- | --- | --- |
|  | SPEED | C4 | BITON | CGP | C7 | C6 | ACSN | CP | C5 | C3 |  |  |
| Informative signatures | 6 | 270 | 11 | 233 | 185 | 1 | 8 | 150 | 64 | 9 | 15 | 10 |
| Total | 11 | 858 | 40 | 1698 | 2436 | 99 | 63 | 1330 | 1454 | 615 | 50 | 326 |
| Fraction | **55%** | **31%** | **28%** | 14% | 8% | 1% | **13%** | 11% | 4% | 1% | **30%** | 3% |

### Informative gene sets tend to be much more frequently significant in some typical cancer data analysis

Our hypothesis that an informative gene set has much higher chances to be enriched in a typical transcriptomic data analysis, is here tested according to the procedure detailed in the Methods section in three typical scenarios: (i) KRAS mutated vs. wild type colorectal cancer ^10^; (ii) metastatic vs. primary colon cancer ^11^ and (iii) tumor vs. normal in four tissues (Lung ^12^, Gastric ^13^, Colon ^14^, Cervix ^15^). As shown in Table 2, in all three cases the number of informative signatures in the GSEA output was strongly enriched (average P-value 10^-79). Note that, while the selection of the informative signature was performed using an unsupervised approach, the validations presented in this section are realized using a supervised one (GSEA). Nevertheless the amount of informative signatures obtained in the output of the GSEA analysis is significantly higher than what could be expected at random.

**Table 2.**
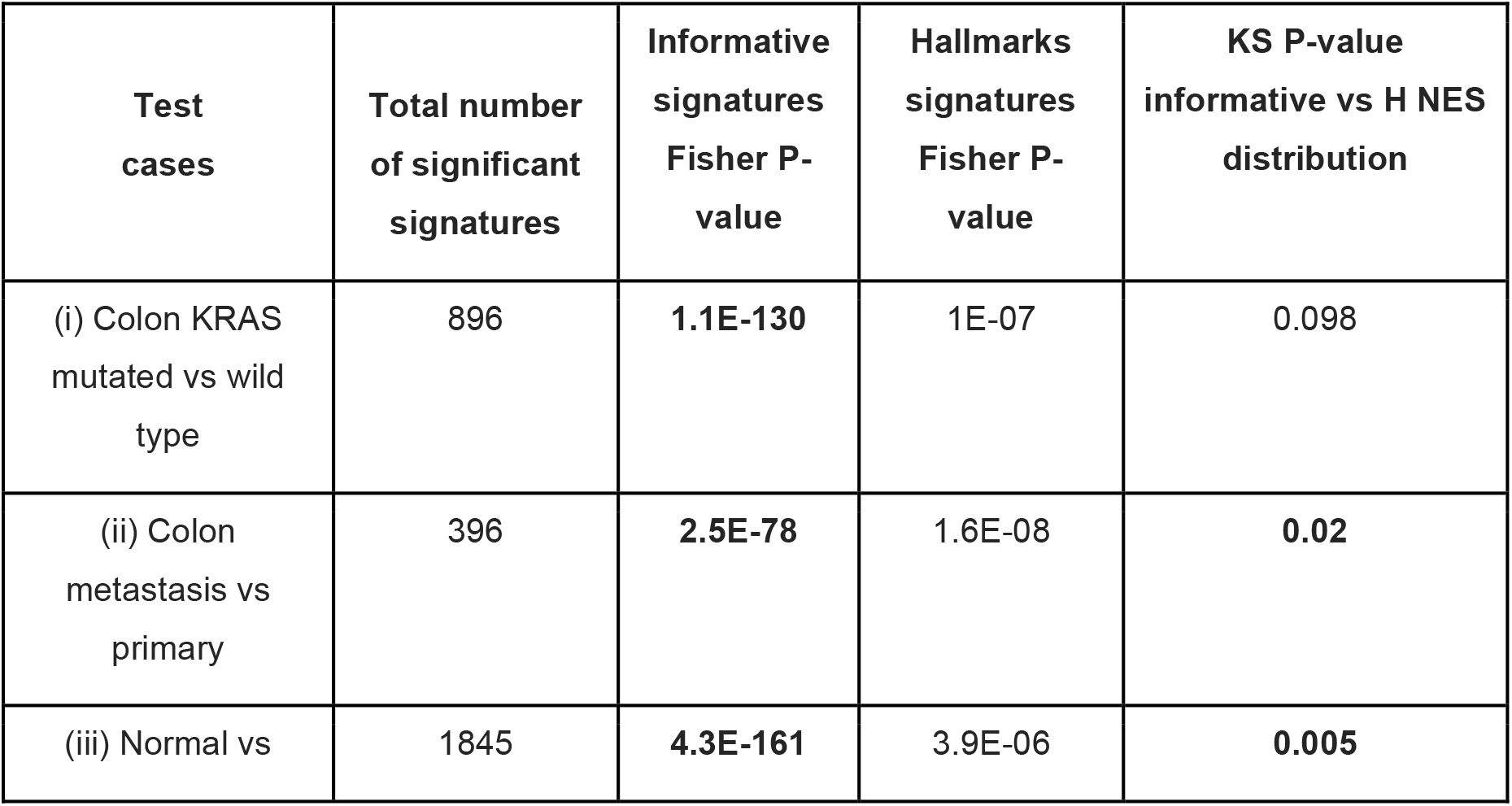
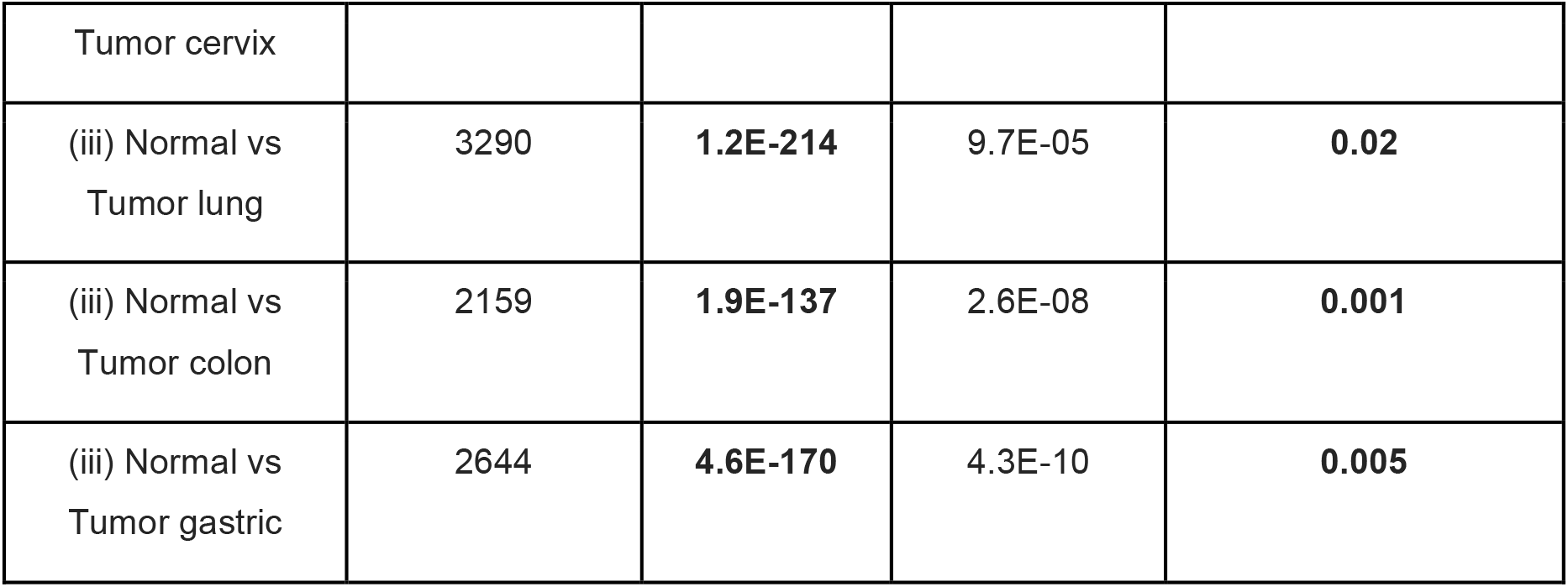
Performances of the informative and Hallmark collection in respect to the full set of starting signatures in three test cases. The columns report for each case: the total number of signatures significant in the test, the Fisher P-value for the informative signatures, the Fisher P-value for the Hallmarks and the Kolmogorov-Smirnov P-value comparing the NES distribution of the informative vs Hallmarks.

### Informative signatures perform better than the MSigDB Hallmarks in some typical cancer data analysis

Given that the only other attempt to prioritize the most reliable non-redundant signatures is represented by the MSigDB Hallmarks, its performances were compared with those of our compendium in the three above test cases in terms of Fisher’s exact test P-values and P-value of the Kolmogorov-Smirnov (KS) test for the NES distributions, as described in the Methods section. The results of both Fisher’s exact test and the KS test are summarized in Table 2. As shown in the table, the Hallmarks signatures obtained significant Fisher P-values in all cases (average P-value 10^-05), confirming the reliability of the procedure employed by Liberzon A. and coauthors. However, the P-values obtained by the Hallmark collection resulted to be always less significant than those of our informative signatures (10^-05 vs 10^-79). Concerning the NES distribution, as shown in Supplementary Figure S1, the informative signatures tend to be always associated with absolute NES higher than those of the Hallmarks. Indeed, the KS P-values are always lower than 0.05, except for the KRAS mutated vs. wild type colorectal cancer example in which the P-value is 0.098. Therefore, not only the informative signatures have higher chances to be significant in a GSEA analysis, but also in the GSEA output they tend to be among those with the highest NES score. This result indicates that our compendium is capturing the strongest sources of expression variation in all three transcriptomic datasets. As a further check, given that 15 over 50 Hallmarks signatures are also contained in our informative collection, the fraction of Hallmarks signatures present in the output of the GSEA analysis and also contained in our informative compendium is evaluated: (i) KRAS mutated vs. wild type colorectal cancer 67%; (ii) metastatic vs. primary colon cancer 67% and (iii) normal tissue vs. tumor in 4 tissues (Lung 48%, Gastric 47%, Colon 44%, Cervix 60%). Such a result shows that among the 50 signatures constituting the MSigDB Hallmarks, those that are found significant in the GSEA analysis are in the majority of the cases also informative. This last observation thus further confirms the reliability of our selection procedure taking into account that the subset of the Hallmarks that we selected tends to be enriched more frequently in the analyzed cancer-specific applications.

### The majority of the informative signatures eigengenes are conserved across cancer types

To further investigate the reliability of our informative collection, we verified if the informative signatures were quantitatively reproduced across different cancer types, where the term quantitatively refers to the eigengenes (set of gene weights) resulting from the PCA decomposition. Computing the conservation score as described in the Methods section and employing a threshold of 10^−6^, 1459 over the 12096 starting signatures (12%) resulted to be conserved across-cancer. On the opposite 703 over the 962 informative signatures (73%) were found to be conserved, showing that the signatures selected with our approach have higher chances to maintain the same quantitative definition across different cancer types and thus they tend to be more robust than the starting ones. Testing then the much stringent threshold of 10^−10^, while the total number of conserved signatures substantially decreased (from 1459 to 408) the percentage of conserved signatures that were found to be also informative substantially increased (from 48% to 83%). This last result thus proves that the previously obtained results are not affected by the value of the chosen conservation score threshold and, more importantly, that the informative signatures are always among the most across-cancer conserved gene sets.

### Functional redundancy of gene sets is poorly explained by their intersection size

Two gene sets with empty intersection can represent incomplete transcriptional read-outs of the same biological process (i.e., cell cycle), being thus functionally redundant. For example, when data-derived expression signatures are constructed, the number of genes whose expression is associated with the phenotype of interest is generally minimized and only the most representative are maintained in the signature. This procedure may lead to the reconstruction of two data-derived signatures associated to the same phenotype but having a poor/null intersection. This leads to a well-known problem of signature reproducibility ^16^. In order to quantify the scale of this phenomenon, the two redundancy measures: functional redundancy, computed as described in the Methods section, and gene content intersection in terms of Jaccard-index (JI) are compared in Figure 2B. As expected, high JI value usually results in high functional redundancy (i.e. high average correlation between meta-samples over all cancer types). However, as shown in the figure, high functional redundancy is distributed over a large range of JI values with a surprisingly higher points density in the area corresponding to poorly overlapping gene sets. Therefore, in order to reduce the functional redundancy between gene sets, it is not sufficient to simply take into account their overlap, but also their correlation of activity needs to be considered. A first consequence of this result is that the Hallmarks collection, based on the use of the JI as a measure of redundancy, is not able to completely capture analogous signatures. To better quantify redundancy, also the meta-samples correlation need to be considered. The intrinsic limitation of such approach is that it requires the wide employment of expression data and thus its output is data-dependent.

### InfoSigMap a user-friendly interactive representation of the functional redundancy structure between informative signatures for insightful gene set score visualization

Given the predefined collection of informative signatures, GSEA or alternative approaches can be employed in order to score them based on a transcriptomics dataset and a set of sample labels. The output of such analysis consists of a table of gene sets enrichment statistics. This tabular organization of the output, containing functionally connected signatures scattered throughout, frequently does not help the interpretation of the results and the formulation of consistent biological hypothesis. In order to improve this aspect, we developed InfoSigMap (http://navicell.curie.fr/pages/maps_avcorrmodulenet.html), according to the procedure described in Methods. As shown in Figure 3, the obtained network contains six main strongly connected components of informative signatures. The first is associated to core cellular functions (i.e. all those basic functions that are fundamental for the life of the cell) and it contains cell cycle, mRNA translation, splicing, MYC targets, protein degradation and oxidative phosphorylation. Of particular interest is the fact that in this component the network is able to clearly separate the signatures associated to the different cell cycle phases. The second connected component is instead related to the tumor microenvironment and it comprises: immune system, inflammation, TNF-α pathway, interferon and extracellular matrix/EMT. The third and fourth connected components, smaller than the others, correspond to transcription and neuronal system. Finally, the fifth and sixth components contain signatures associated to analyses performed on the same expression compendia (GNF2 ^17^ and GCM ^18^) and simply represent genes with large neighbourhood overlap. The first and the second large components are connected through an area associated to Experimental perturbations of Immune cells (EI) signatures. Indeed the informative signatures derived from EI (dark red nodes in Figure 3) are separated into two main areas: one, as could be expected, belongs to the tumor microenvironment component and it is strongly linked to the immune system/inflammation signatures; the second, instead, is part of the proliferation component, strongly linked to the cell cycle area. We considered that such unexpected configuration could be caused by the presence of gene sets, belonging to the EI category, but obtained from differentiation induction experiments or on immortalized cell lines, and thus characterized by differences in the proliferation rate. Indeed an alteration in the cell cycle process could justify the high correlation of activity (reflected in an high density of links) present between these two sets of signatures. In order to test this hypothesis, a dataset obtained from the expression profiling of human CD4+ T cell during differentiation induction was employed ^19^. The informative pathways altered during the experiment (differentiated vs undifferentiated) were detected employing ROMA according to the procedure described in Methods section. As shown in Supplementary Figure S2, the two areas of experimental perturbations of immune cells signatures have an opposite behavior concordant with that of the signatures around them. Indeed the area near to the cell cycle results to be donwregulated in the differentiated cells, while those that are part of the immune island are upregulated. This result confirms our starting hypothesis that an alteration of the cell cycle process was at the origin of the observed subdivision and it suggests that a reshaping of the EI category would be recommended for their future use in data analysis. Another non-intuitive observation is that signatures coming from the same collection tend to co-localize in the map and data-derived signatures tend to be clearly separated from knowledge-based ones. The discrepancy between data-derived and knowledge-based signatures can be explained by the fact that the transcriptional readouts of a biological process might be very different from the genes involved in the process itself. Yet another unexpected observation is that higher functional redundancy exists between signatures of the same collection rather than between signatures describing the same biological function, suggesting that all the analyzed signature collections can be prone to a common bias. The only two exceptions to this trend to some extent are the MSigDB Hallmark collection and the SPEED signatures (although several non-informative SPEED signatures are clustered together). These compendia indeed resulted to be well spread around the map, confirming that they are able to well capture the main biological signals encoded in the transcriptomic data. Nevertheless, some areas such as mRNA translation, transcription, splicing and protein degradation were not covered by any of the Hallmark and SPEED signatures, indicating that there is still the need of other signatures in order to have a complete portrait of the transcriptomic landscape. As introduced above, InfoSigMap was developed to simplify the navigation and interpretation of the gene set score distributions. In the next section, some examples of typical analysis scenarios where InfoSigMap can be employed to formulate consistent biological hypothesis are presented.

**Figure 3.**
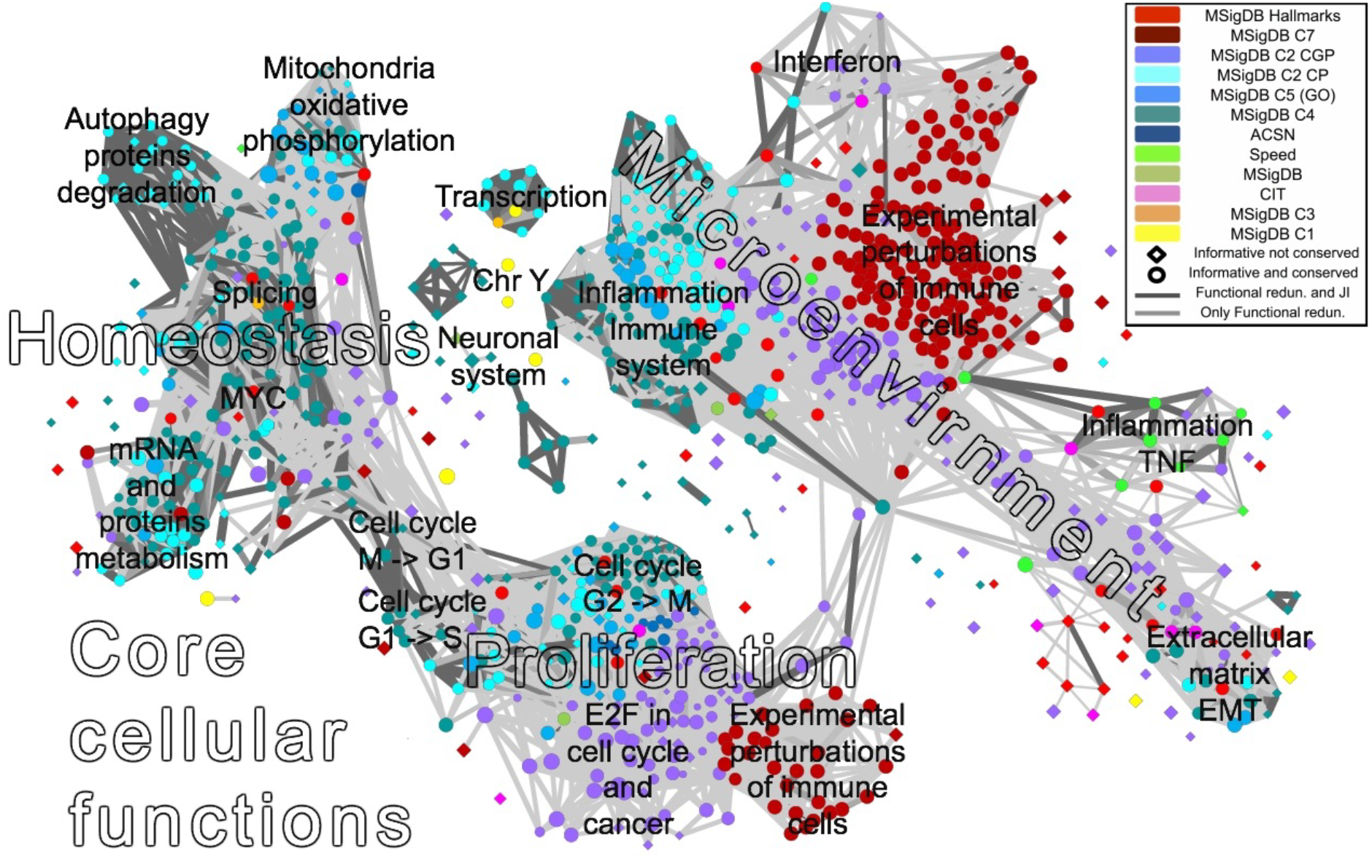
InfoSigMap: user-friendly interactive representation of the informative signatures. The network map of the 962 informative signatures plus SPEED and Hallmarks is here reported as available on the website (http://navicell.curie.fr/pages/maps_avcorrmodulenet.html). The signatures are organized as nodes of the network. Nodes colors correspond to the different signatures categories, while the shape is a diamond for informative signatures and circular for those that are also conserved. The links correspond to redundancy between couples of gene sets (functional redundancy in light gray, dark gray if also the Jaccard-index intersection is significant). The names annotated on the top of the map denote areas of the network containing signatures associated to same biological function. The interactive on-line version of this map can be browsed as an instance of Google Maps, with the possibility of zooming in and /out, getting description of gene signatures, and visualizing data (various gene set scores, such as GSEA or ROMA scores) on top of the map.

### InfoSigMap can be used to visualize the results of transcriptomics data analysis

InfoSigMap is tested to investigate the alterations affecting the transcriptome of the three aforementioned typical cancer problems: (i) KRAS mutated vs. wild type colorectal cancer; (ii) metastatic vs. primary colon cancer and (iii) tumor vs. normal tissue in lung, gastric, colon and cervix. In all three cases, the procedure employing ROMA and described in Methods is applied resulting in the output shown in Figure 4. Below the obtained results are discussed in detail:

**Figure 4.**
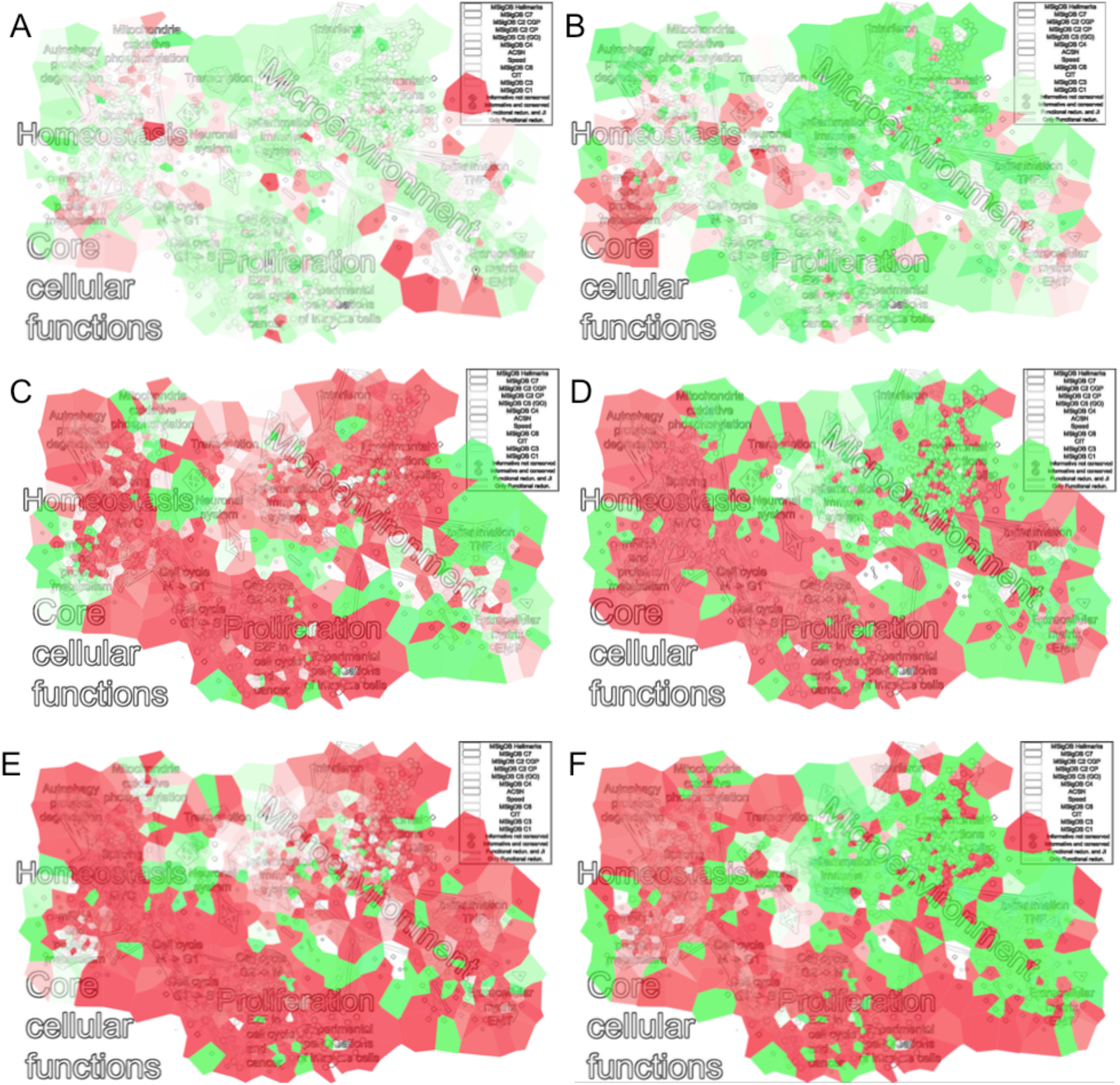
Results of InfoSigMap applied to some typical data analysis scenarios. Four examples showing how InfoSigMap provides a more insightful interpretation of the lists of significant signatures output of the classical gene expression analysis tools is here proposed. The significant fold-changes resulting from the differential ROMA analysis are plotted on the top of the map according to a heatmap coloring highlighting up-(red) and down-regulated (green) gene signatures. The plots are organized as follows : (a) KRAS mutated vs. wild type colorectal cancer; (b) metastatic vs. primary colon cancer and (c-f) tumor vs. normal tissue in cervix, colon, gastric and lung, respectively.

#### (i) KRAS mutated vs. wild type colorectal cancer

The impact of *KRAS* mutation on CRC transcriptome is investigated with InfoSigMap (Figure 4A). *KRAS* mutated CRC patients are known to be resistant to standard Epidermal Growth Factor Receptor (*EGFR*) inhibitory treatments ^20,21^. The output of our analysis can thus give some indications concerning possible new processes to be targeted in KRAS mutated patients. The strongest effect reported in Figure 4A (bright red area) is the upregulation of a subset of the metastatic signatures. This result fits with previous evidences that KRAS mutation is associated to metastasis in patients with CRC ^22,23^. Moreover an alteration of the metabolism is detectable from an upregulation of the mitochondria and oxidative phosphorylation area. This result fits with previous experimental evidences. Indeed *KRAS* mutation has already been shown to induce mitochondrial oxidative stress, inducing a phenotype consistent with the so-called Warburg effect, a metabolic alteration fundamental for cancer cell proliferation ^24–26^. In CRC, *KRAS* mutation also causes an alteration of the transcriptional response and amino acid metabolism machineries, two fundamental processes for cancer cell proliferation and maintenance ^27,28^. This effect is captured in our analysis by the upregulation of the mRNA translation/protein metabolism area on the InfoSigMap network.

#### (ii) Metastatic vs. primary colon cancer

The differential module activity between metastatic and primary Colon Cancer (CC) is studied (Figure 4B). As expected, an up-regulation of the collagen/EMT area of the network clearly appears on the map Figure 4B. Among the signatures found in this area, we can observe an upregulation of the miR-21 targets whose role in EMT is well-known ^29,30^. Moreover, the network areas of mRNA and proteins metabolism and splicing resulted to be significantly upregulated. This is not surprising given that the aberration of the RNA processing machinery (stability, metabolism, splicing and polyadenylation) is known to be associated with cancer initiation and progression. In CC, beta-catenin (*CTNNB1*), involved in the Wnt pathway, is generally the cause of the RNA processing alterations ^31–33^. This is confirmed in InfoSigMap; indeed *CTNNB1* is found active as shown by the upregulation of its targets (node FEVR_CTNNB1_TARGETS). The study of the cancer-specific RNA metabolism is a relatively unexplored area of research, with potentially significant implications for the prevention and treatment of CC. The above results confirm the experimentally observed *CTNNB1*-mediated alteration of the RNA processing machinery. On the other side, a strong downregulation of the cells proliferative activity can be observed. This phenomenon has already been documented and found associated with poor-prognosis in colorectal cancer (CRC) ^34,35^. The observed slow-proliferation in metastatic CC may be caused by a high proportion of cancer stem-like cells. Indeed stem cells are in a quiescent state, a phenomenon that could explain the cell cycle downregulation detected in our analysis. The hypothesis of a high stem cell concentration is also confirmed by the significant downregulation of the immune area. Indeed the stem-like phenotype of metastasis-initiating cells is generally associated with immune evasive quiescence, even if this point is not well documented in CC ^36^.

#### (iii) Tumor vs. normal in four tissues

When then compare tumor vs normal tissue in cervix, colon, gastric and lung cancer (Figure 4C-F). A global feature present in all four tissue types is the upregulation of the connected component associated to the core cellular functions. This is not a surprising result, given that cancer cells generally inactivate tumor suppressors and hyperactivate oncogenes to promote sustained proliferation, alter autophagy and the various steps of the RNA transcription and translation processing machinery, develop metabolic imbalances and enhance resistance to mitochondrial apoptosis ^37^. The microenvironment-associated connected component instead shows a dual behavior. It is indeed significantly upregulated in cervical and gastric cancer and downregulated in colon and lung. The results thus suggest a different role of the immune system in these four tumors. A possible explanation for this result is that the tumors are associated to different levels of antigenicity, i.e. the extent to which tumor cells display HLA-restricted antigens that can be selectively or specifically recognized by T cells ^38^. Tumors with low antigenicity hide against cytotoxic attack leading to a passive escape from anti-cancer immune defense. This hypothesis is supported by the observation that the HLA signature present in our network (GNF2_HLA_C) is concordantly downregulated in lung and colon and upregulated in cervix and gastric. Moreover, concerning lung cancer, its association with low antigenicity had already been observed ^39^. The tumor antigenicity is one of the aspects that seem to determine if a patient will respond to a given immunotherapy. In this sense, a comprehensive pan-cancer classification of the immune component behavior could give indications regarding those individuals who are most likely to respond to immune-based therapies.

## DISCUSSION

Data-driven and a priori-knowledge-based gene signatures are largely employed in cancer studies to score clinical samples according to distinct tumor subtypes, identify important cellular responses to stimuli, predict clinical outcomes and quantify the activation of signaling pathways. Nowadays signature collections are getting larger providing the benefit of a more complete coverage of the existing biological processes. However, the growth of these compendia is posing two main challenges due to the reliability and redundancy of the collected gene sets.

Here, we developed a new methodology for assessing the value of a gene set which is based on the notion of informative signature, i.e. a gene set able to systematically robustly rank tumor samples in many independent datasets. A restricted collection of 962 informative gene sets is suggested for transcriptomic data analysis in cancer biology. The robustness of the information content enclosed in our compendium is then proved showing that an informative gene set has much higher chances to be selected (enriched) in a typical scenario of transcriptomic data analyses, even in the ones using supervised methods, and that the eigengenes of the majority of the informative signatures tend to be conserved across cancer types. The redundancy of the informative collection is then investigated, showing that functional redundancy is a frequent phenomenon not captured by the approaches previously proposed for gene signatures summarization. To integrate all the obtained results we developed InfoSigMap, a user-friendly interface designed for insightful data visualization. InfoSigMap was applied in some typical scenarios of cancer data analysis. The obtained results proved that a global view of the concordant behavior of functionally redundant signatures leads to an insightful interpretation of the results in respect to what can be deduced from the lists of significant signatures output of gene expression data analysis tools. In all four analyzed cases, the obtained results were found to fit with the previous experimental knowledge, confirming the reliability of our approach. However, also some indications concerning new candidate mechanisms to be experimentally investigated were extracted, showing how InfoSigMap can help the formulation of new biological hypothesis.

## METHODS

### Definition of “an informative signature”

To explain the rationale behind the definition of informative signature, let us consider to apply Principal Component Analysis (PCA) to a gene expression data table whose columns correspond to the genes from a selected gene set and whose rows correspond to samples. If we observe that the variance explained by the first principal component computed for such a table is significantly larger than for a random set of genes of the same size then the considered gene set is called overdispersed ^40,41^. Intuitively, an overdispersed gene set has a stronger contribution to the data variance than expected by chance. Similarly, if the ratio between the variances explained by the first and second principal components computed for the aforementioned table is larger than for a random set of genes then the given gene set is called coordinated. Intuitively, the existence of a statistically significant gap between the first and the second eigenvalue of the covariance matrix corresponds to an overall increase in the pairwise correlations between the genes of the signature compared to what can be observed at random. The advantage of having a coordinated gene set is that it defines an axis of principal variance in the multi-dimensional distribution of samples and thus it robustly ranks samples independently on their labeling, as discussed in the Introduction (see Figure 1). In the context of cancer biology, we define informative a gene set that is simultaneously overdispersed and coordinated in more than two cancer types.

### Transcriptomics data input of the analysis

To systematically search for informative signatures, a large pan-cancer TCGA compendium of gene expression data derived from 32 solid cancer types (ACC, BLCA, BRCA, CESC, CHOL, COAD, DLBC, ESCA, GBM, HNSC, KICH, KIRC, KIRP, LGG, LIHC, LUAD, LUSC, MESO, OV, PAAD, PCPG, PRAD, READ, SARC, SKCM, STAD, TGCT, THCA, THYM, UCEC, UCS, UVM) was employed. The data were downloaded from TCGA and normalized. An overview of the samples available for the different tumor types is reported in Supplementary Table S1.

### Signatures collection input of our analysis

A vast collection composed of both data-derived and *a priori*-knowledge-based signatures was considered as input for our analysis. The signature collections: Molecular Signature Database (MsigDB v5.2) ^42^, Atlas of Cancer Signaling Network (ACSN) ^43^, the top-contributing genes of the components identified by Biton et al. (here denoted as CIT) ^44^ and the Signaling Pathway Enrichment using Experimental Data sets (SPEED) ^8^ have been downloaded and organized, obtaining a starting collection of 12096 signatures. In the following, we will consider as data-derived signatures: CIT, SPEED and some MSigDB categories (clusters of genes co-expressed in microarray compendia (C4), signatures of oncogenic pathway activation (C6), the large collection of immunological conditions (C7) and chemical and genetic perturbations (CGP) part of the MSigDB collection canonical pathways and experimental signatures curated from publications (C2)). On the other side, ACSN and the MSigDB categories: genes sharing cis-regulatory motifs up- or downstream of their coding sequence (C3), genes grouped according to Gene Ontology (GO) categories (C5) and Canonical Pathways (CP), part of C2 and including the well-known BIOCARTA, KEGG and REACTOME databases, will be denoted as knowledge-based. Finally, the MSigDB collections genes grouped by their location in the human genome (C1) and the Hallmarks (H) will not be associated to any of the two previous classifications.

### Procedure for the prioritization of those signatures that are informative in cancer biology

To detect which of the starting 12096 signatures, detailed above, were informative, we employed the Representation and quantification Of Module Activity (ROMA) tool, designed for the robust detection of overdispersed and coordinated modules ^41^. The activity of each signature was thus evaluated in all the 32, previously described, expression datasets separately. Only those signatures having the p-values associated to the variance explained by the first principal component (ROMA L1 score) and to the ratio between the variances explained by the first and second principal components (ROMA L1/L2 score) lower than 0.05 in at least two tumor datasets were prioritized.

### Test if informative gene sets are more frequently enriched in typical cancer analysis scenarios

Our hypothesis that an informative gene set has much higher chances to be enriched in a typical transcriptomic data analysis, is tested by using Gene Set Enrichement Analysis (GSEA), a well-known and widely adopted supervised approach ^42^. GSEA was applied to three typical cancer-related problems using the entire collection of 12096 starting signatures. The set of significant signatures was determined selecting those with a GSEA FDR q-value lower than 0.05. Then, the enrichment of our set of informative signatures in the output of the GSEA analysis was evaluated through a Fisher’s exact test.

### Comparison Informative signatures vs Hallmarks in typical cancer analysis scenarios

Given that only one other attempt to prioritize the most reliable non-redundant signatures exists and it is represented by the MSigDB Hallmarks, the procedure described in the previous section was repeated also for this database. The performances of our informative collection were then compared with those of the Hallmarks in terms of Fisher’s exact test P-values. Moreover, the distributions of the absolute GSEA Normalized Enrichment Score (NES) for the two collections were studied and the significance of the difference between the two distributions was evaluated through a Kolmogorov-Smirnov (KS) test.

### Evaluation of the signatures eigengenes conservation across-cancers

To further investigate the reliability of our informative collection, we verified if the informative signatures were quantitatively reproduced across different cancer types, where the term quantitatively refers to the eigengenes (set of gene weights) resulting from the PCA decomposition. For each informative signature, the pair-wise correlation between the eigengenes obtained in the 32 cancer types were computed and a conservation score was obtained as the geometric mean of the Pearson correlation p-values. We define a conserved gene set by having the conservation score lower than 10^−6^. To then evaluate how much the results of this test were affected by the threshold used to define a conserved gene set, also a much stringent threshold of 10^−10^ was tested.

### Comparison between functional redundancy and intersection size of gene sets

Two gene sets with empty intersection can represent incomplete transcriptional read-outs of the same biological process, being thus functionally redundant. In order to quantify the scale of this phenomenon, we have compared the normalized size of gene set intersection with their functional redundancy measure. For each couple of informative signatures, the average correlation between their meta-samples was thus computed over the 32 cancer types. This approach captures all those couples of gene sets having a similar pan-cancer behavior and thus representing the same transcriptional read-out, independently on whether they have significant number of common genes or not. To evaluate if the functional redundancy between a couple of signatures is well explained by their overlap size, the intersection in terms of Jaccard-index (JI) between all couples of informative gene sets was computed and the two measures were compared.

### InfoSigMap construction procedure

To help the quick navigation of the set of informative signatures and to improve the interpretation of their concordant behavior, we developed InfoSigMap (http://navicell.curie.fr/pages/maps_avcorrmodulenet.html), a user-friendly Google-Maps based visualization method. The construction of InfoSigMap involved three main steps: (i) creation of the signature redundancy graph; (ii) definition of its layout and (iii) representation of the graph as an interactive online map. In line with what has been already done in Enrichment Map, GOIorize and ClueGO ^2,4,6^, the first step is performed by organizing the 990 signatures (corresponding to the 962 informative collection plus all Hallmarks and SPEED signatures even if they were not shown to be informative) into a weighted network, where each signature is a node and links represent redundancy between couples of gene sets. Differently from the previously mentioned Cytoscape plug-ins, the links of our network are weighted averaging over two measures of signatures redundancy: overlap (JI) and functional redundancy. The functional redundancy was computed as the average correlation coefficients between the metasamples defined by ROMA, for each pair of informative signatures in each cancer type. The signatures having average correlation above 0.7 are connected in the graph and the final weights of the links were obtained as the mean between the average correlation and the JI. The threshold 0.7 is justified by appearance of distinguishable but still connected functional components in the graph. For the second step of graph structure representation, a different shape is used to denote the gene sets that are only informative (diamond) and those that are also conserved (circle), while the node size denotes the number of genes in the signature. Links are also classified into two classes, dark gray is used for those edges that connect signatures being both functionally redundant and having a significant JI, while light gray denotes links only associated to functional redundancy. Finally, the thickness of the links is proportional to their weights, the standard Cytoscape organic layout is used to spatially organize the largest connected component of the network and smaller components or unconnected nodes were positioned by using the structure of weaker correlations. The areas of the network containing signatures associated to same biological functions are then identified and manually annotated on the top of the map to help the navigation of the users. In addition, also a purely data-driven layout was computed by applying tSNE dimension reduction method to the matrix of average pairwise correlations between the meta-samples defined by our signatures in all cancer types. This view of the InfoSigMap is available at <u>http://navicell.curie.fr/pages/maps_avcorrmodulenet.html</u> (View/tSNE selection in the right-hand panel). Finally, the representation of the network as an interactive online map is achieved by using NaviCell ^45^, powered by Google Maps API.

### Using InfoSigMap to have a global view of the signatures behavior

Gene sets can be tested for differential activity across different experimental conditions using a tool of choice (e.g. GSEA, ROMA). If ROMA is chosen for this test, first the activity of the informative signatures is evaluated by applying ROMA, then the differential module activity is evaluated by Student’s t-test and fold-change applied to the ROMA activity scores. Finally, the fold-changes associated to a significant Student’s t-test P-value (lower than 0.05) are mapped to the nodes of InfoSigMap as a color gradient, from red (up-regulation) to white (no significant change) to green (down-regulation), using the map staining approach described in ^45^. The map is thus colored in the territories around each node creating a continuous colored pattern that helps a qualitative appreciation of the concordant/discordant behavior of large map regions.

## AUTHOR CONTRIBUTIONS

LCan, AZ and LCal designed the study. LCan made numerical computations. LCan, AZ wrote the manuscript. NB, MR provided materials and analyzed the results. LCan, LM participated in case studies. All authors have read and edited the manuscript.

## ACKNOWLEDGMENTS

This study was financially supported by COLOSYS EU EracoSysMed project.

## Supplementary Information Pan-cancer classification of gene signatures for their information value and functional redundancy

### Supplementary contents

**Table S1.** Number of samples available for each transcriptomic dataset used for the analysis

| **tumor code** | **tumor name extended** | **number of samples** |
| --- | --- | --- |
| ACC | Adrenocortical carcinoma | 78 |
| BLCA | Bladder Urothelial Carcinoma | 408 |
| BRCA | Breast invasive carcinoma | 1085 |
| CESC | Cervical squamous cell carcinoma and endocervical adenocarcinoma | 303 |
| CHOL | Cholangiocarcinoma | 36 |
| COAD | Colon adenocarcinoma | 276 |
| DLBC | Lymphoid Neoplasm Diffuse Large B-cell Lymphoma | 48 |
| ESCA | Esophageal carcinoma | 182 |
| GBM | Glioblastoma multiforme | 155 |
| HNSC | Head and Neck squamous cell carcinoma | 515 |
| KICH | Kidney Chromophobe | 65 |
| KIRC | Kidney renal clear cell carcinoma | 516 |
| KIRP | Kidney renal papillary cell carcinoma | 286 |
| LGG | Brain Lower Grade Glioma | 513 |
| LIHC | Liver hepatocellular carcinoma | 368 |
| LUAD | Lung adenocarcinoma | 510 |
| LUSC | Lung squamous cell carcinoma | 488 |
| MESO | Mesothelioma | 87 |
| OV | Ovarian serous cystadenocarcinoma | 302 |
| PAAD | Pancreatic adenocarcinoma | 177 |
| PCPG | Pheochromocytoma and Paraganglioma | 181 |
| PRAD | Prostate adenocarcinoma | 494 |
| READ | Rectum adenocarcinoma | 90 |
| SARC | Sarcoma | 255 |
| SKCM | Skin Cutaneous Melanoma | 103 |
| STAD | Stomach adenocarcinoma | 401 |
| TGCT | Testicular Germ Cell Tumors | 138 |
| THCA | Thyroid carcinoma | 501 |
| THYM | Thymoma | 120 |
| UCEC | Uterine Corpus Endometrial Carcinoma | 173 |
| UCS | Uterine Carcinosarcoma | 57 |
| UVM | Uveal Melanoma | 80 |

**Table S2.**
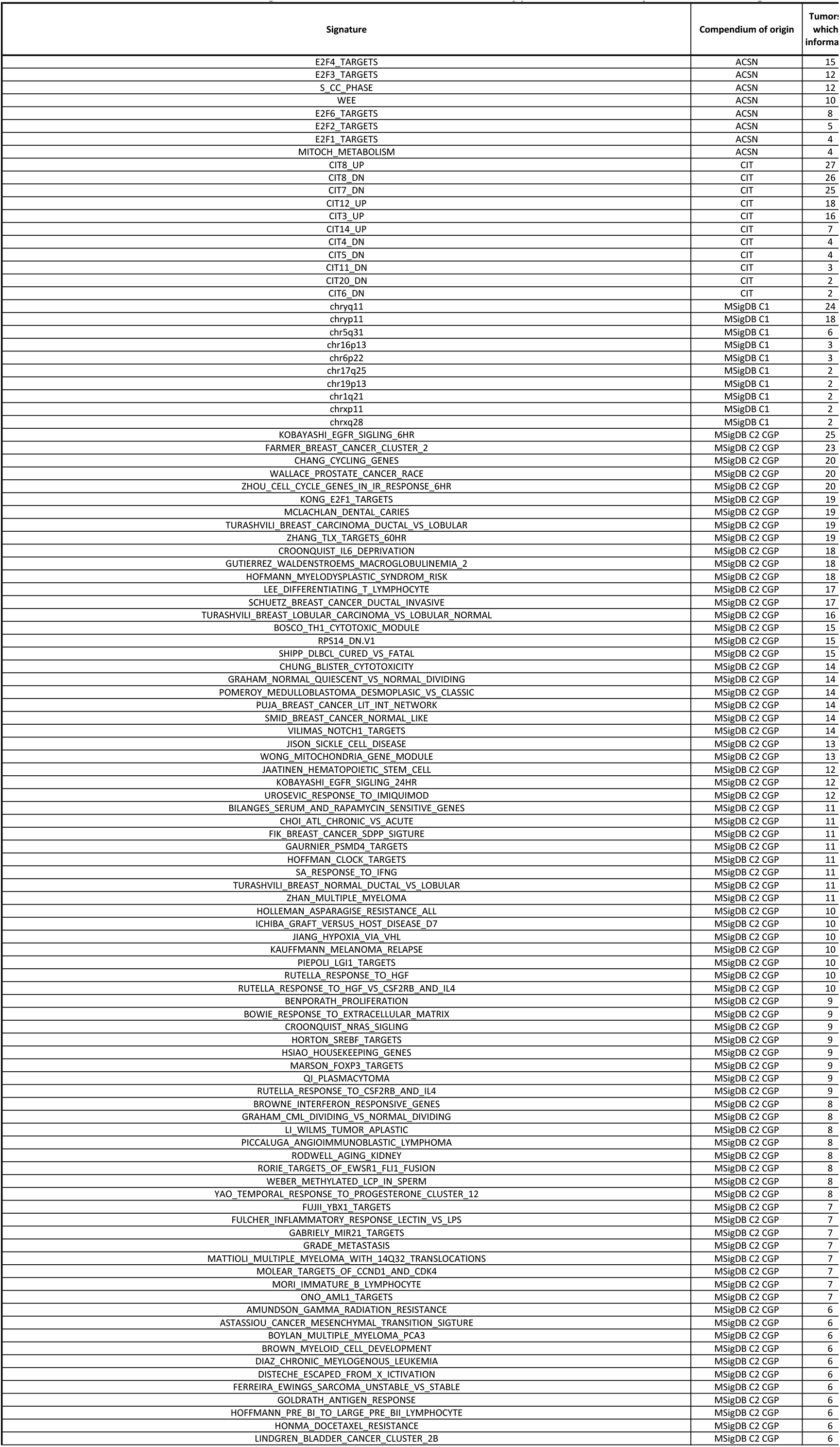
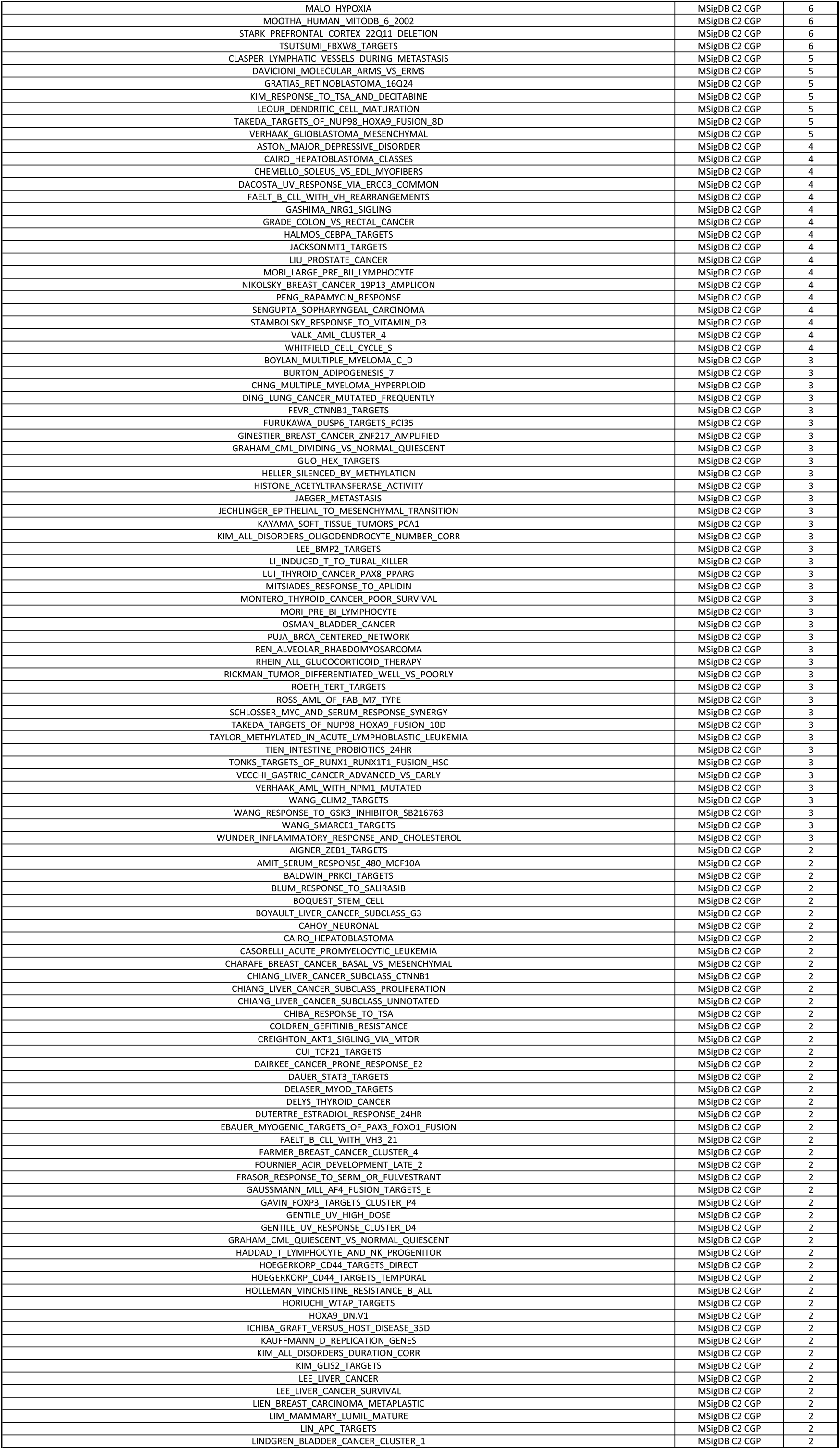
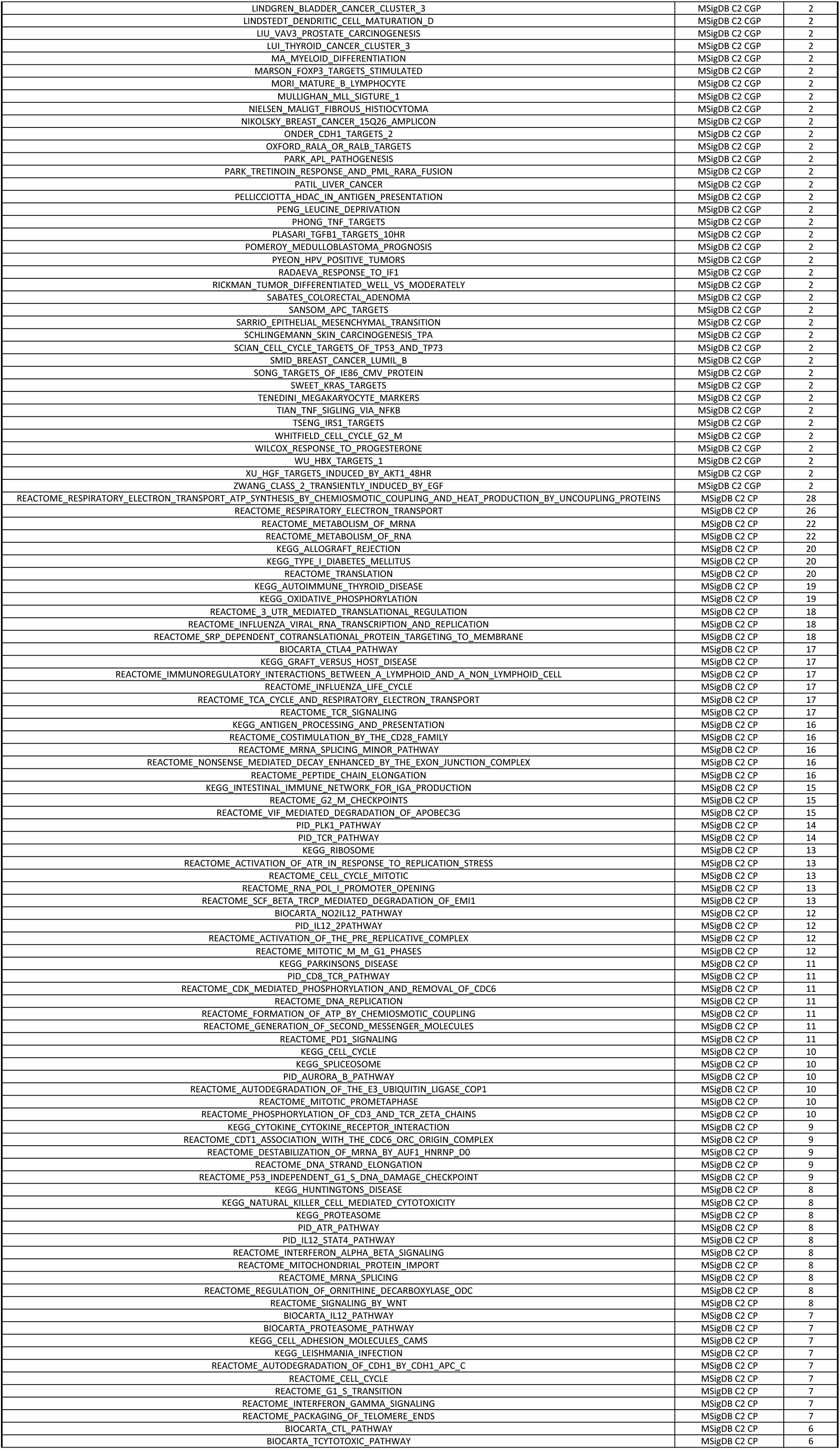
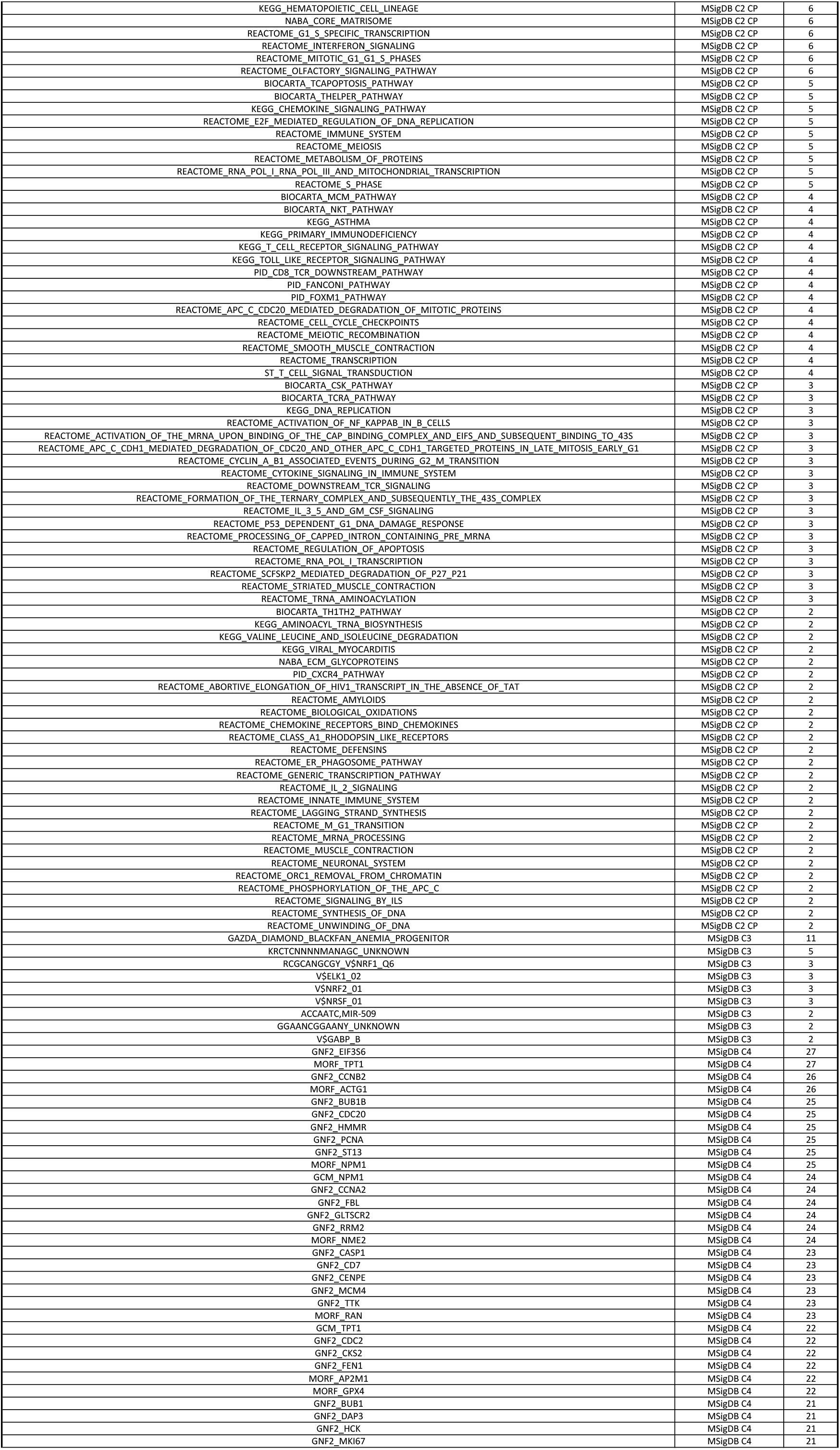
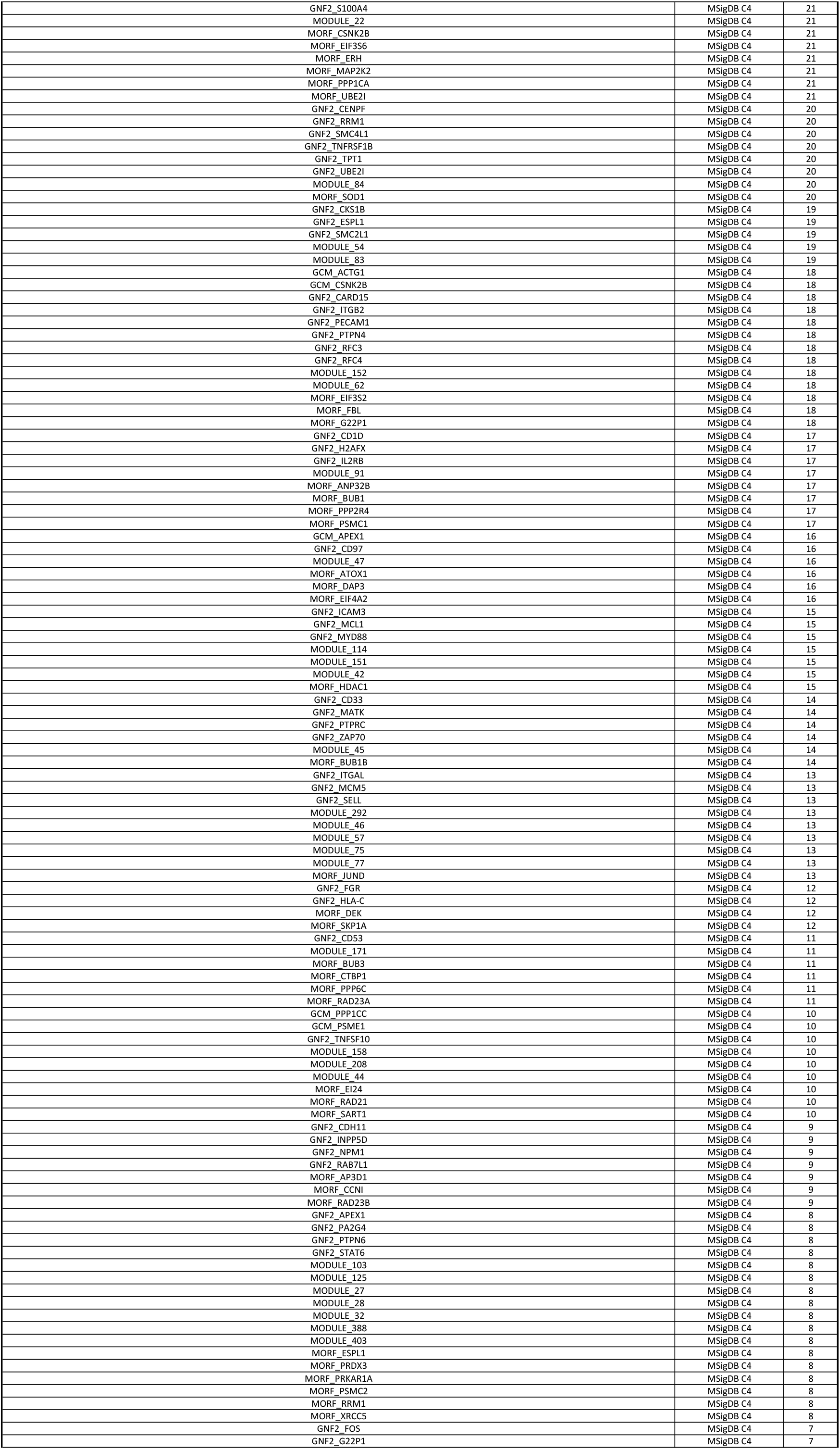
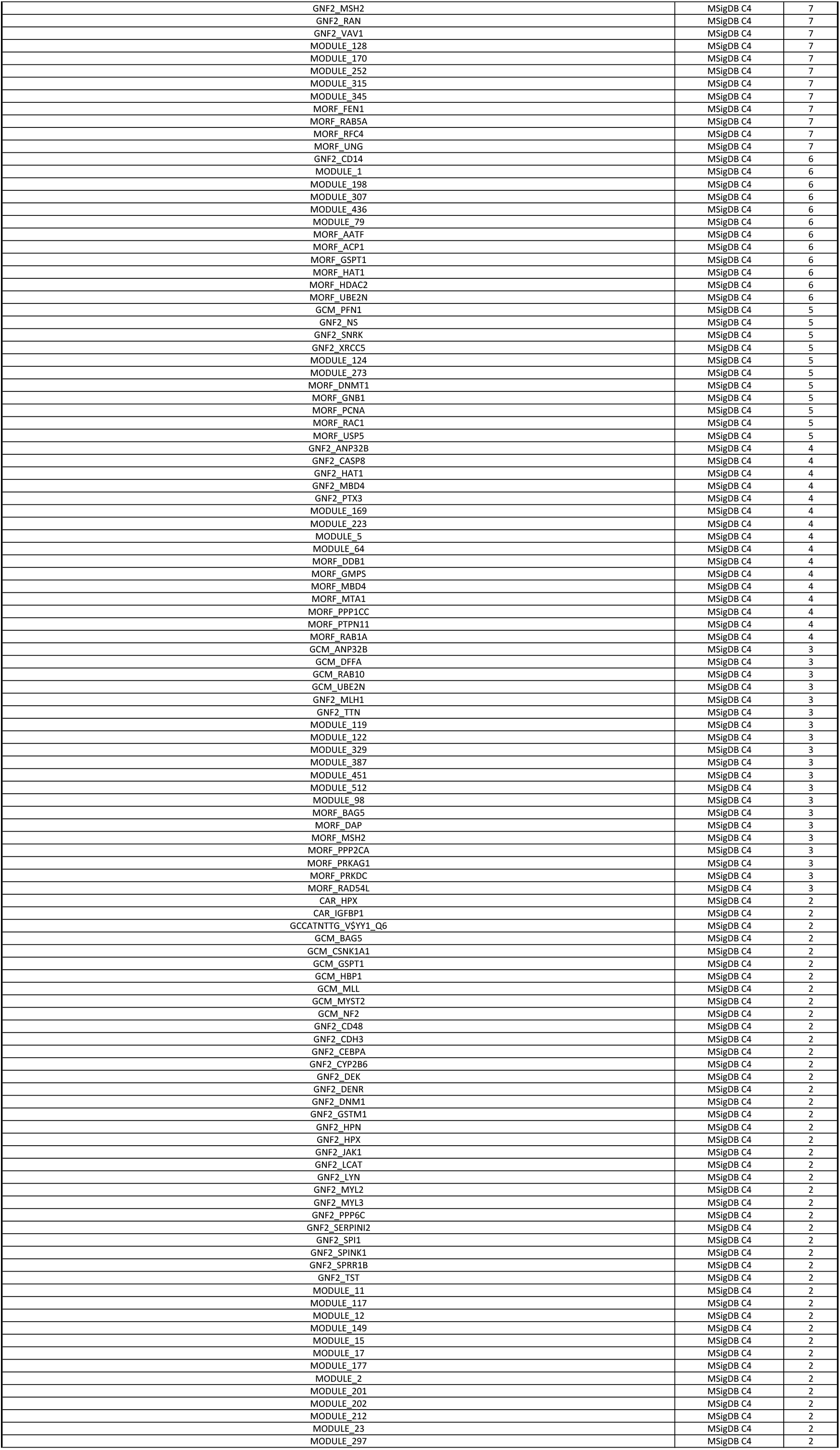
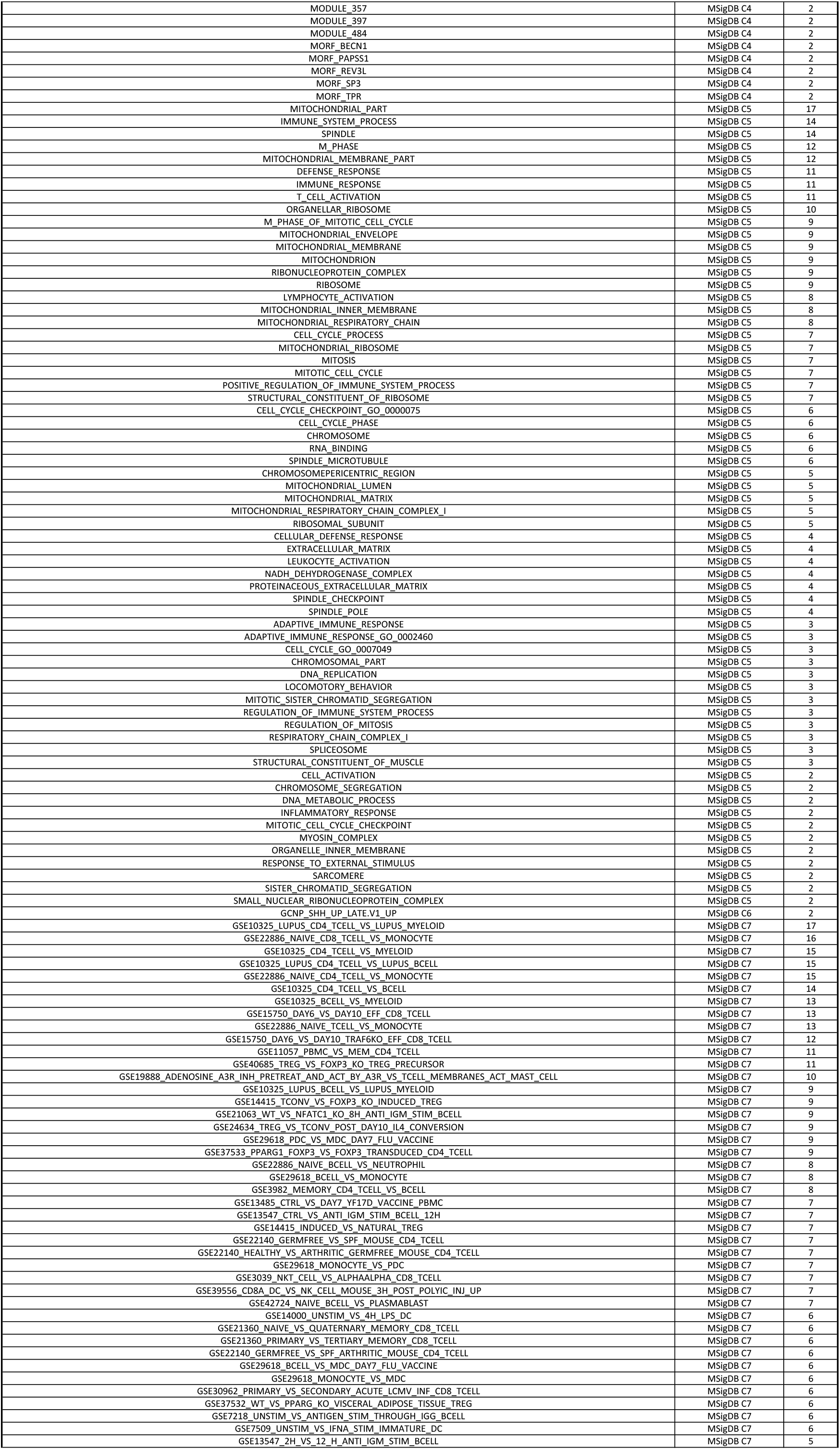
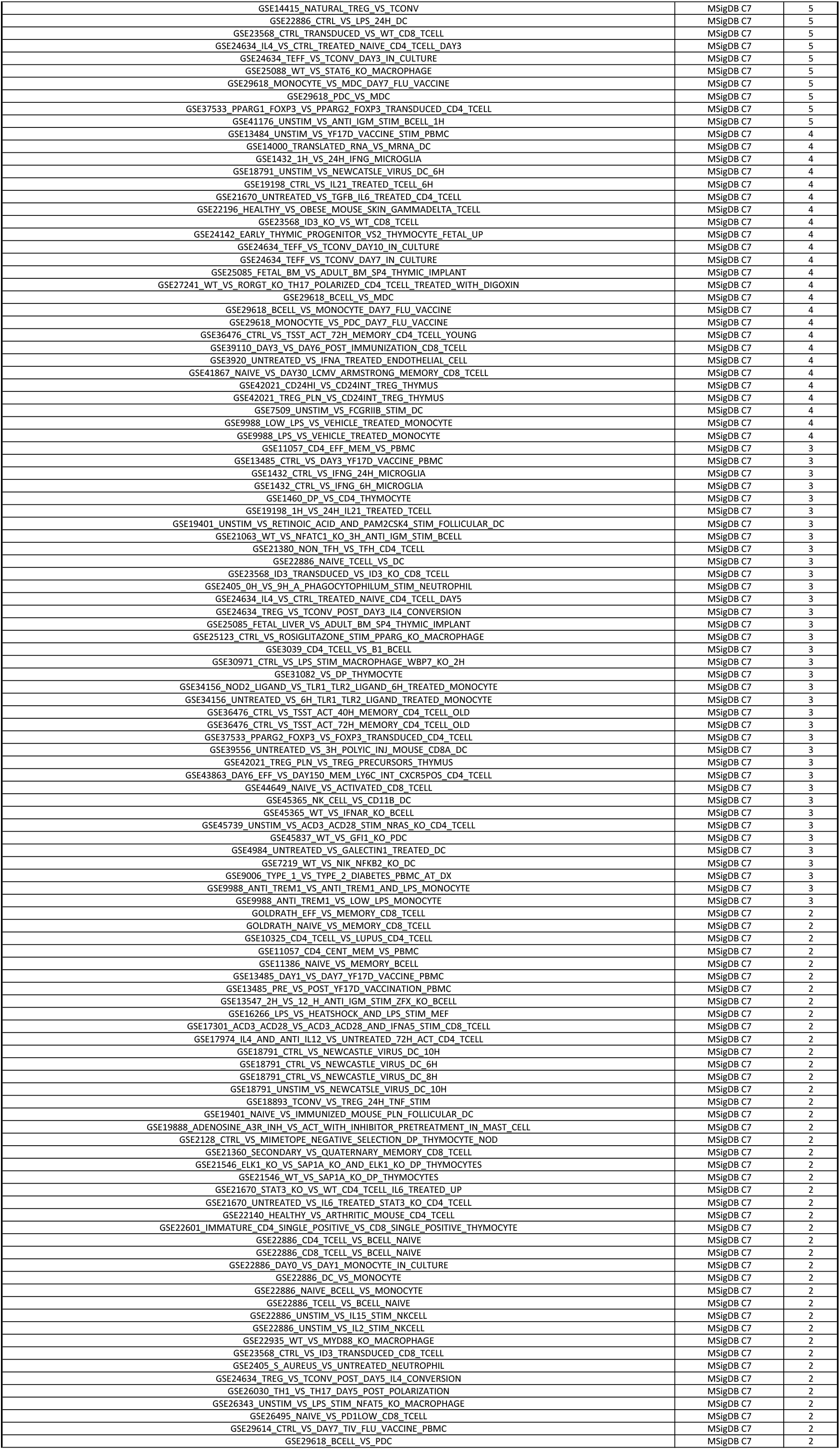
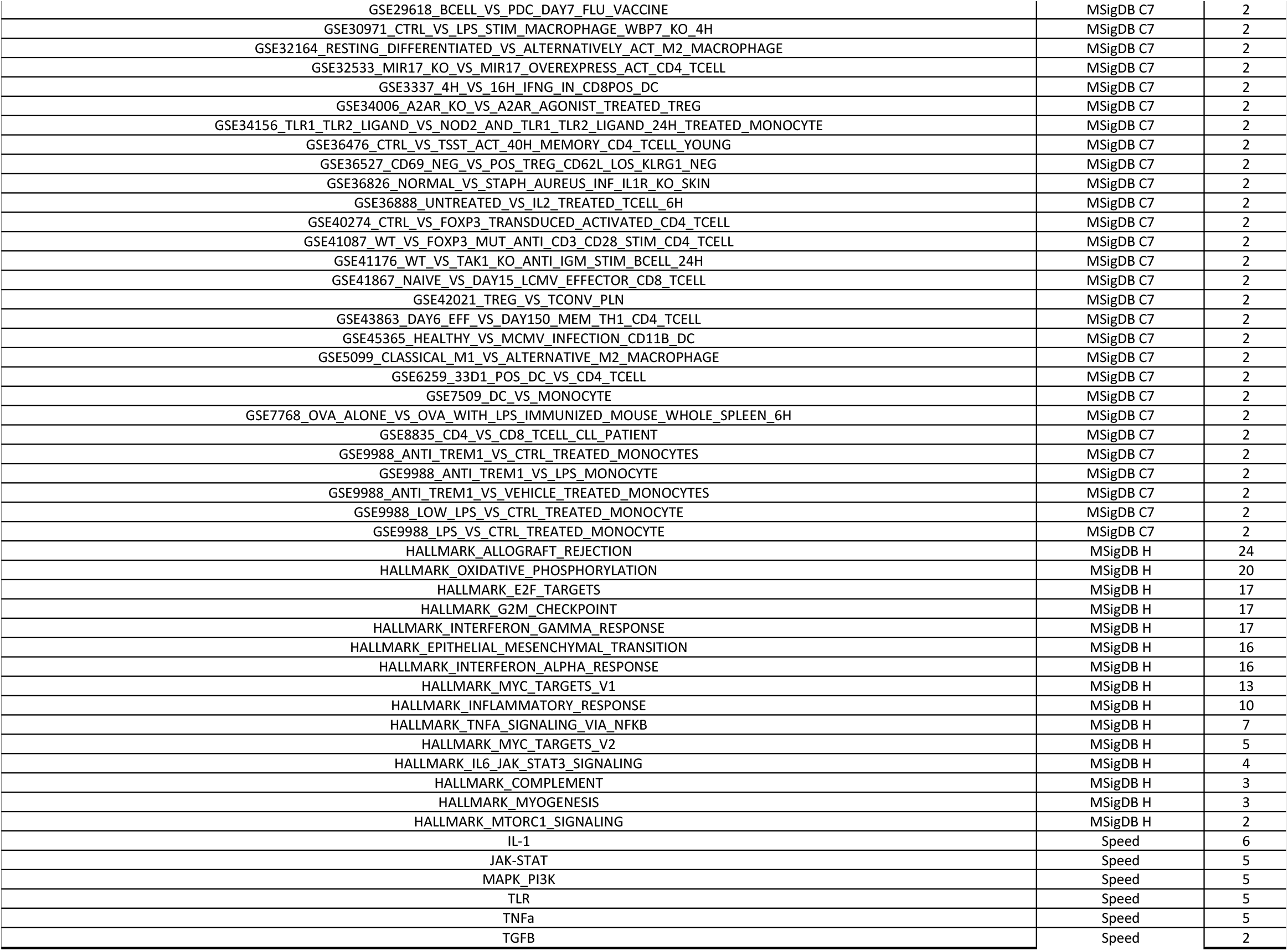
List of informative signatures and number of tumors in which they were found significant

**Figure S1.**
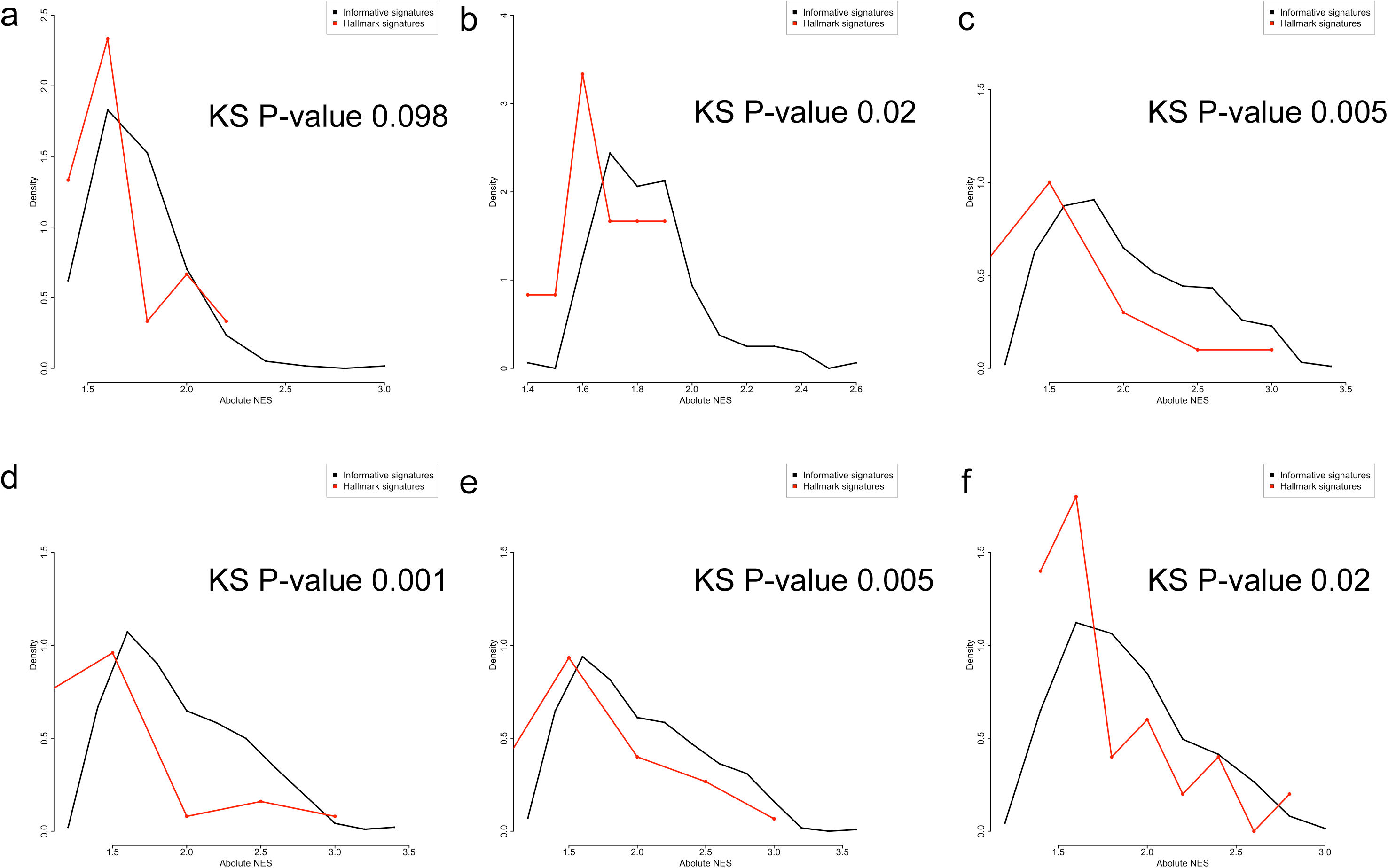
Comparison GSEA absolute NES values for the informative GSEA significant (black) vs. Hallmark GSEA significant (red) in (a) KRAS mutated vs. wild type colorectal cancer; (b) metastatic vs. primary colon cancer and (c-f) tumor vs. normal in cervix, colon, gastric and lung, respectively.

**Figure S2.**
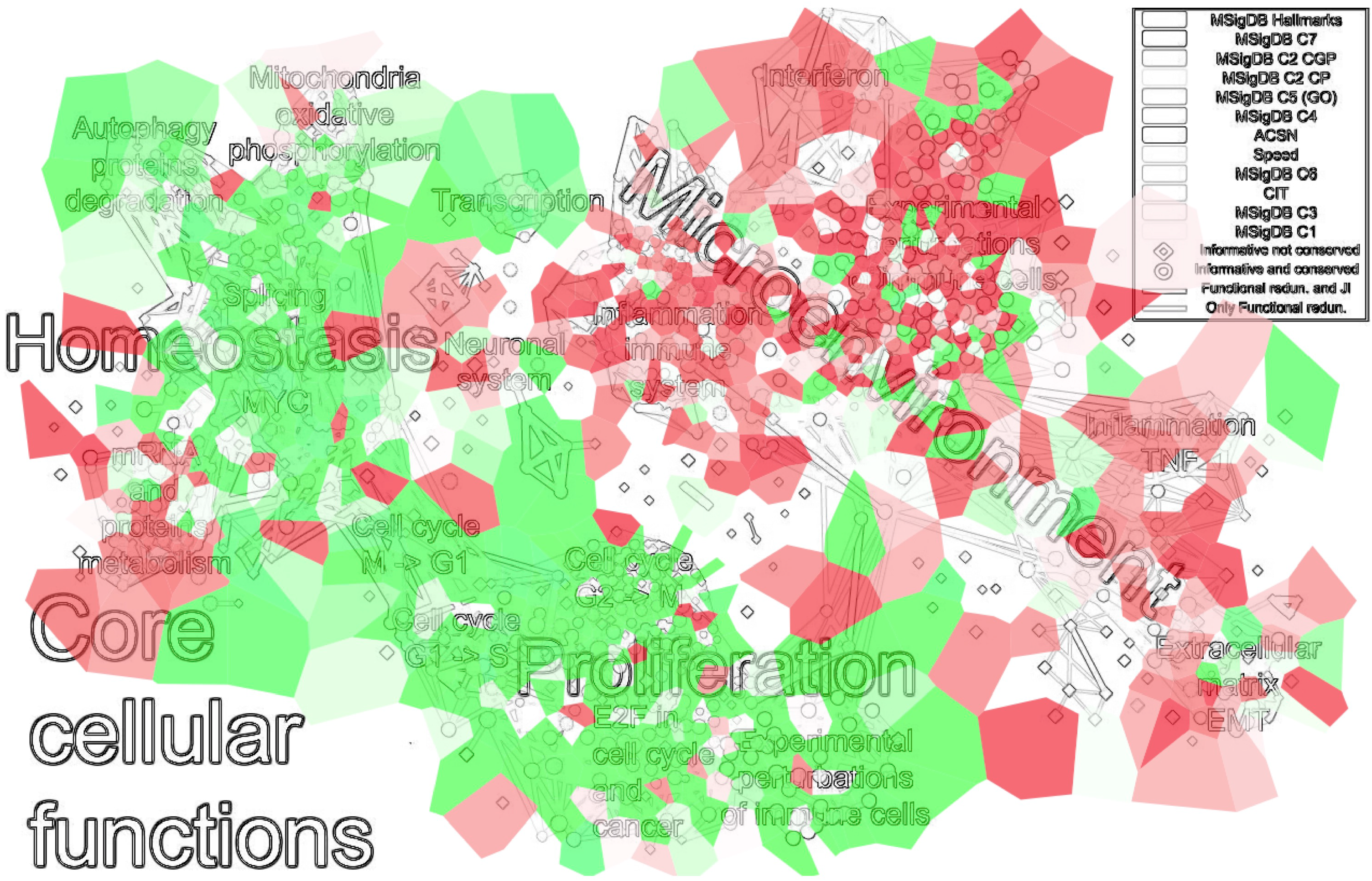
Fold changes of ROMA activity on the expression profiling of human CD4+ T cell during differentiation induction plotted on InfoSigMap. Red, and green indicate upregulation and downregulation, respectively.

